# Cell villages and Dirichlet modeling map human cell fitness genetics

**DOI:** 10.1101/2025.09.26.678880

**Authors:** Chloe Hanson, Tim Derebenskiy, Ana Rodriguez Vega, Yashika S. Kamte, Rachel G. Fox, Hannah Lambing, Laila Sathe, Tyler E. Dietterich, Patrick Allard, Harold Pimentel, Michael F. Wells

## Abstract

The capacity of cells to proliferate and survive is central to development and disease. Assays that measure cell fitness are therefore a cornerstone of biology, but traditional techniques lack donor diversity and have high technical variability that impedes scale and reproducibility. To overcome these barriers, we designed and validated a “cell village”-based fitness screening approach using pooled cultures of 12–39 genetically distinct human neural progenitor cell (NPC) lines. We also developed Townlet to establish a foundational statistical framework based on Dirichlet regression for analyzing proportional data from cell villages. Applying these systems, we identified hyperproliferation in NPCs harboring the autism risk factor chromosome 16p11.2 deletion, mapped common genetic variants near *ZFHX3* associated with NPC proliferation rate, and discovered genetic modifiers of lead (Pb) sensitivity implicating *ARNT2*. Together, these experimental and analytical tools advance a scalable, genetically diverse *in vitro* platform for dissecting human variation in cell fitness and gene-environment interactions.

## Introduction

Cell proliferation and maintenance are fundamental to life. During early development, cells increase in number over time and differentiate to build tissues and organs through processes governed by genetics and the environment. In humans, differences across individuals in proliferation and viability mechanisms (i.e., cell fitness) contribute to variability in complex traits, such as organ size, regeneration capacity, and resilience to extrinsic stressors ^1–4^. Likewise, between-individual variation in the penetrance of deleterious genetic mutations or the susceptibility to harmful environmental exposures can drive differential risk for developmental disorders, tissue degeneration, and cancer ^5–8^. Accurate measurements of cell fitness that take into account numerous genetic backgrounds and environmental stimuli are therefore essential for elucidating the mechanisms that coordinate cell proliferation and viability under healthy and disease states, as well as for the screening and development of effective therapeutics.

Human pluripotent stem cells (hPSCs) and their derivatives are valuable tools for studying fitness in genetically distinct cells. Conventional hPSC-based approaches utilize an arrayed culture format, in which cells from different individuals (i.e., donors) are plated into separate wells for experimentation and analysis. These designs, however, are often limited by low throughput, small donor sample size, and high well-to-well and batch-to-batch technical variation that can mask biological signals and inhibit reproducibility ^9–14^.

To increase the scale, genetic diversity, and number of perturbations in a single experiment’s design, we and others recently introduced “cell villages,” which consist of tens to hundreds of genetically distinct human cell lines cultured together in shared *in vitro* environments ^15–18^. This strategy enables quantification of cel-lular phenotypes across many genetic backgrounds in parallel under uniform culture conditions and reduced technical variation. Villages have been deployed to interrogate between-donor variation in differentiation potential ^19,20^, viral susceptibility ^15^, and drug response ^17^. However, villages have not been systematically used to quantify cell proliferation or viability, nor have computational models been developed that consider the unique design and structure of these experiments.

Here, we establish experimental and analytical tools for village-based fitness assays of human neural progenitor cells (NPCs), which are the building blocks of the fetal brain. Many genetic and environmental risk factors for neurodevelopmental disorders alter NPC proliferation and viability ^21–24^. We constructed multiple hPSC-derived NPC villages consisting of 12-39 donors (Tables S1-S2) to quantify and explain between-donor variation in proliferation and sensitivity to environmental stress. Given that the inherent compositional ^25^ (i.e., proportional) data structure of villages invalidates the use of traditional statistical tools like ANOVA, we built a user-friendly hierarchical Dirichlet regression model called Townlet that analyzes village donor proportion data from low coverage DNA sequencing ^17^ collected over multiple time points and/or perturbations to estimate individual donor fitness.

Using Townlet, we tested several biological hypotheses and addressed common questions about village experimental design, power, reproducibility, and the impact of cell non-autonomous factors. We first devised a statistical simulation framework to evaluate our model’s improved performance over more classical statis-tical models (e.g., linear regression) and to assist in designing future high-powered, reproducible cell village experiments. We then demonstrated how our model can answer biologically relevant questions by: (i) showing that donors harboring neurodevelopmental disorder genetic risk factor chromosome 16p11.2 deletion dis-play NPC hyperproliferation, (ii) deciphering how common genetic variation contributes to between-donor differences in NPC proliferation rates, and (iii) nominating genetic modifiers of sensitivity to the known neu-rodevelopmental toxicant lead (Pb). Our integrated experimental format elucidates natural human variation in cell fitness while revealing how gene-environment interactions amplify or buffer these effects.

## Results

### Villages are reproducible and capture cell intrinsic proliferation rates

Though gaining in popularity, important questions remain regarding the cell village approach, such as the extent to which pooling donors minimizes technical variation and the influence of cell non-autonomous factors (i.e., between-donor cellular interactions) on phenotypes. To address these questions and validate villages for *in vitro* cell fitness assays, we measured the NPC proliferation rates of multiple donors cultured in village and array formats and quantified reproducibility across wells, batches, and formats.

First, we pooled hPSC-derived NPCs ^15,26^ from 12 donors and plated them into three technical replicate wells (“Batch 1”) to measure well-to-well reproducibility. We then passaged the replicates independently every 5 days for 15 days, harvesting DNA at each passage. We collected 0.5X-coverage whole genome sequencing data from each sample for Census-seq, a computational tool that infers the donor composition of each replicate (Figure 1A-1B; Data S1) ^17^. Donor proportions were highly correlated between well replicates within a batch (Figure 1C-1D; Spearman and Pearson *r >* 0.996). To test village batch reproducibility, we repeated the experiment on the same 12-donor NPC village constructed 4 days later (“Batch 2”) and found that donor proportion estimates were strongly correlated between Batch 1 and Batch 2 (Figure 1E-1F; *r >* 0.939*, p <* 1.56 · 10*^−^*^17^). Thus, although donor cell proportions evolve over time, their trajectories are consistent both within and between batches.

**Figure 1.**
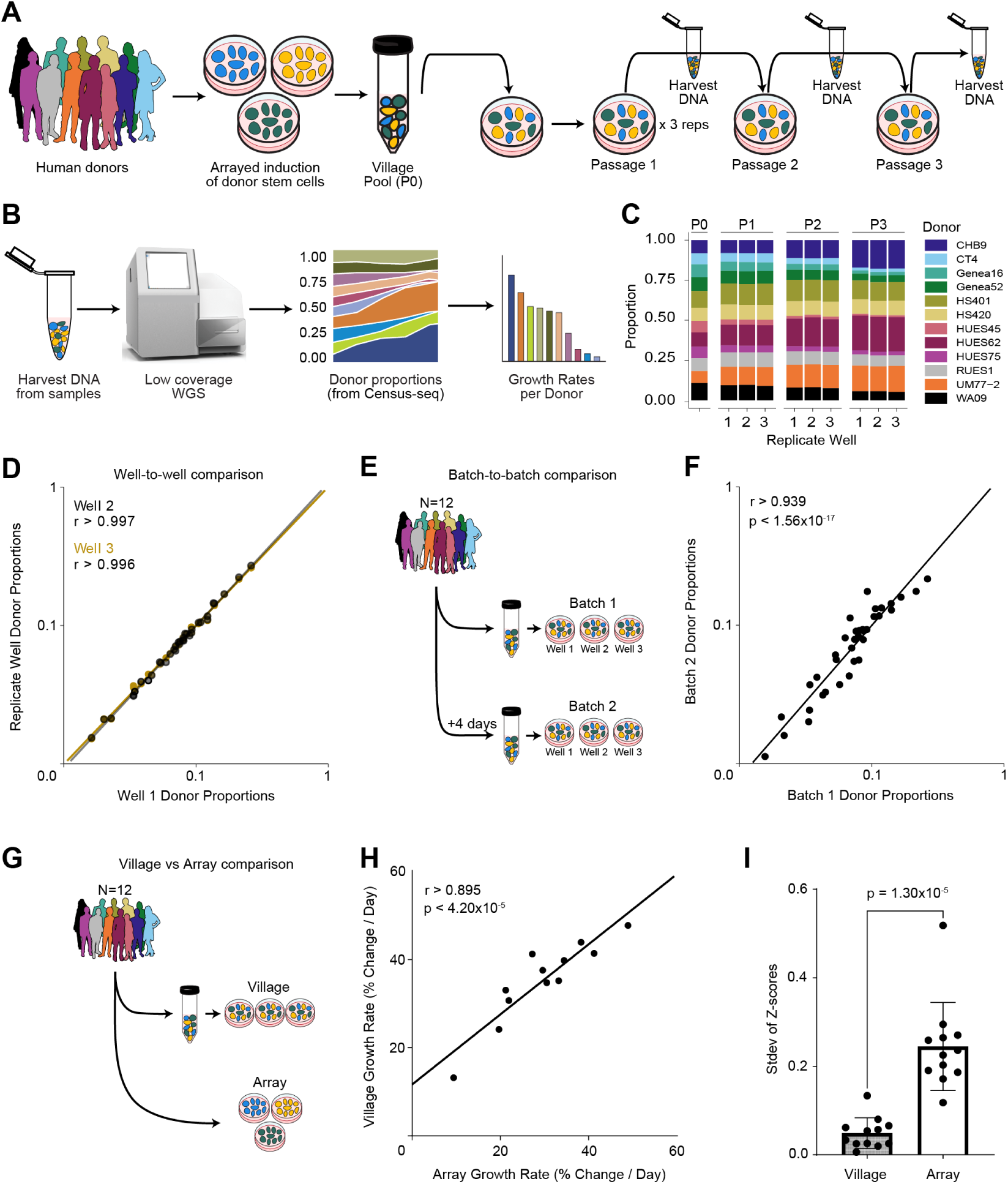
Validation of village-based NPC fitness assay. (A) Schematic of cell village workflow. Human pluripotent stem cells from 12 donors were induced to NPCs separately in an arrayed format prior to pooling (Passage 0; P0) and plating. Village cells were harvested in triplicate for Census-seq at each of the three passages. (B) Census-seq workflow. DNA from frozen cell pellets is sequenced at low (0.5X) coverage. Census-seq estimates the proportion of each donor’s DNA in a sample via expectation-maximization, and growth rates are calculated for each donor based on their changing proportions over time. (C) Census-seq donor proportion estimates from a 12-donor NPC village at P0 and three passages (P1-P3). Individual donor proportions are represented with different colors. (D), Donor proportions from 12-donor NPC village are highly correlated between replicates within a batch (Well 2 vs Well 1, *p* = 3.69 *·* 10*^−^*^39^, Pearson r = 0.998, Spearman r = 0.997; Well 3 vs Well 1, *p* = 1.63 *·* 10*^−^*^37^, Pearson r = 0.997, Spearman r = 0.996) (E) Batch comparison experimental design. A 12 donor NPC village was constructed (Batch 1). Four days later, the same 12 donors were pooled into a village (Batch 2). (F) Donor proportions are strongly correlated between batches. Each data point represents a donor’s mean proportion at a given passage (Pearson r = 0.907, Spearman r = 0.939, *p <* 2.40 *·* 10*^−^*^15^; biological n = 12 donors; technical n = 3 per time point). (G) Village-array comparison. Twelve donor NPC lines were pooled into a village or plated separately in an arrayed format for fluorescent-based cell counting and growth rate analysis. (H) Donor intrinsic growth rates are maintained in a village. Each data point represents a donor’s mean exponential growth rate (continuous, % per day) in both a village and arrayed format. (Pearson r = 0.909, Spearman r = 0.895, *p <* 4.20 *·* 10*^−^*^5^; biological n = 12 donors; technical n = 3 village wells, technical n = 10 wells per donor in the array). (I) Villages reduce within-donor variation. Donor growth rates are normalized (for the village, within each well (n=3 wells). For the array, within each column (n=10 wells/columns per donor)) to a mean of 0 and standard deviation of 1. Each data point represents the standard deviation across replicates of a donor’s normalized growth rates (biological n = 12 donors; Student’s t-test *p* = 1.30 *·* 10*^−^*^5^).

Co-culture with non-self cells (i.e., other donors) may influence a donor’s cellular behaviors through cell non-autonomous mechanisms ^27^. To test the impact of pooled cultures on donor-level proliferation rates, we plated 12 NPC lines in a standard arrayed format as the reference group and compared their proliferation rates to village-based estimates (Figure 1G). Arrayed exponential proliferation rates for each donor were calculated using an imaging-based approach (Methods); village exponential proliferation rates for each donor were computed by multiplying their Census-seq-derived proportions by total village cell counts measured using an automated hemocytometer. Village and array donor proliferation rates were highly correlated (Figure 1H; *r >* 0.895*, p <* 4.20 · 10*^−^*^5^), suggesting that donors retain their cell-intrinsic proliferation dynamics in a village with minimal cell non-autonomous influence. Additionally, donor proliferation rates within a batch varied significantly less between village replicates than between arrayed replicates (*p* = 1.30 · 10*^−^*^5^; Figure 1I). These results imply that villages capture donor-intrinsic proliferation rates as effectively as arrayed formats but with reduced variation, allowing us to study cell fitness in several genetically diverse donors at once.

### Compositional models accurately estimate donor and group effects in villages

When one donor increases its representation in a village, other donors will decrease proportionally, even in the absence of cell non-autonomous biological influences. Donor proportions are measured using Census-seq, which infers a village’s composition from low-coverage whole genome sequencing ^17^. However, compositional (proportion data which sums to one) data strongly violates the donor independence assumption of linear models due to the dependence of each dimension on the sum of the others. As such, we built Townlet, a hierarchical Dirichlet regression model that estimates individual donor and group proliferation effects in villages using compositional data measured by Census-seq over multiple time points (Figure 2A). All effects are relative to a baseline donor (Methods ) as log-odds, with positive values indicating faster proliferation and negative values slower than the baseline. Additionally, within Townlet’s model, we appropriately estimate uncertainty for calibrated statistical testing and implement a replicate-specific linear model for assessing dispersion (noise) over time in experiments.

**Figure 2.**
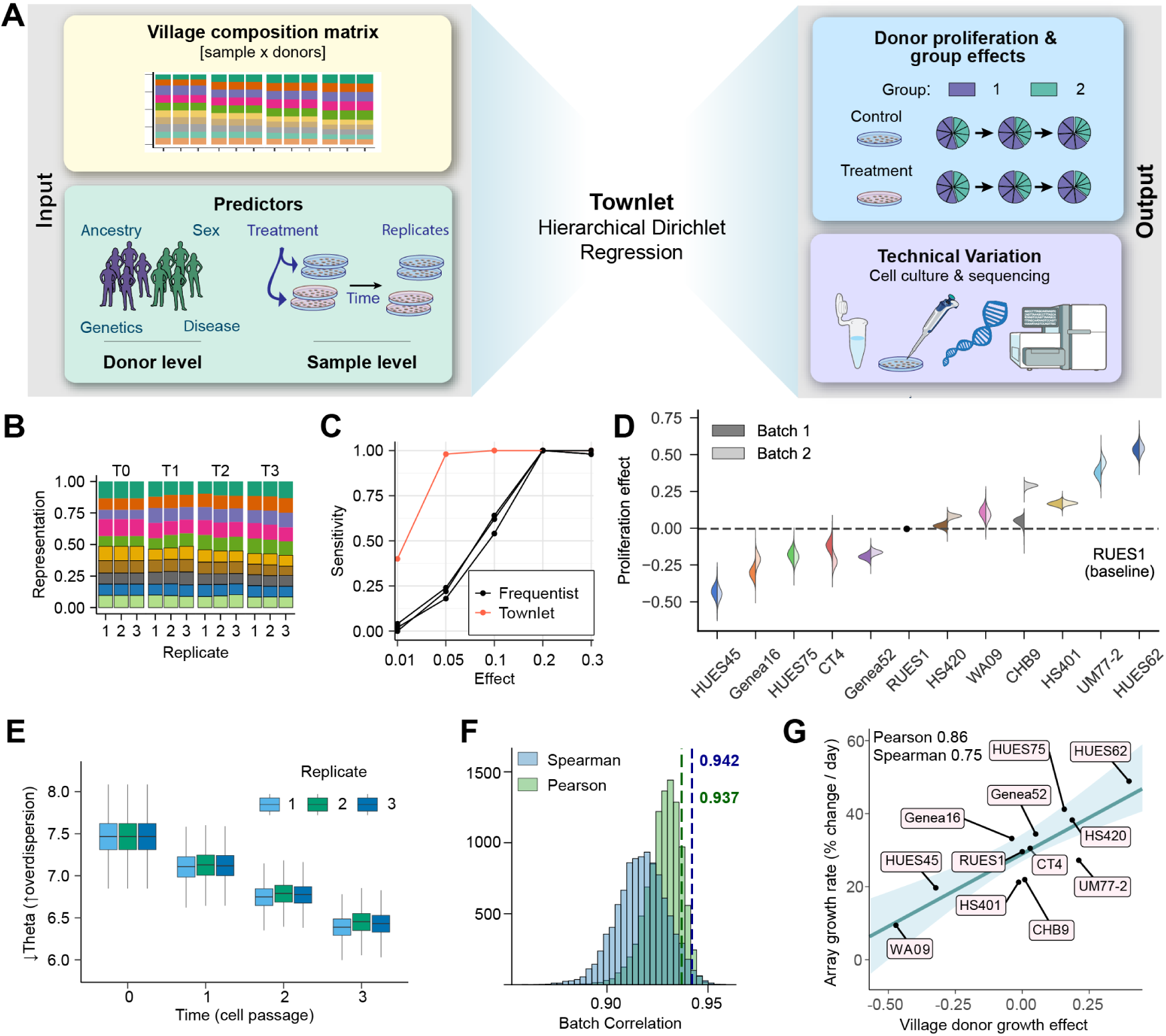
Hierarchical Dirichlet regression for inferring village proliferation and viability. (A) Schematic of Townlet. Inputs include village composition data estimated from the Census-seq pipeline and user defined donor group and sample level predictors. Townlet’s outputs infer significant differences in donor and group proliferation across all treatments and replicate specific technical variation over time. (B) Simulated 10 donor village composition data where all donors have the same initial growth rate, but the donors without the box around them have an additional positive shared group effect that increases their proliferation. (C) Townlet has higher sensitivity compared to other frequentist models even under the simple assumptions outlined in b when simulated group effects are small (*<* 0.2). (D) Proliferation effects estimated from 12 donor villages by Townlet relative to baseline donor RUES1 (dashed line). Split violin shows batch 1 (left) and 2 (right). Donors with distributions above the dashed line grow significantly faster than the baseline, while those falling below grow slower. (E) Overdispersion (technical variation) estimates over time colored by replicate from our 12 donor (Batch 1) experiment. As our model parameter theta decreases, dispersion increases; therefore, in this experiment, replicate technical variation increased consistently over time. (F) Correlation between Batches 1 and 2 donor proliferation effects randomly drawn from posteriors. Dashed lines show high correlation (*>* 0.93) of donor proliferation expectation values. (G) Townlet’s estimated donor proliferation effects correlate strongly with array-based proliferation rates estimated using the exponential growth equation.

We demonstrated the statistical accuracy of Townlet’s framework through simulation and analysis of the 12 donor NPC villages (Batch 1 & 2). Simulation shows that Townlet improves on the standard linear model’s discordant estimates, even when individual donor proliferation differs subtly (Figure 2B-2C). We inferred log-odds donor proliferation effect from each batch (Figure 2D) and modeled experiment dispersion per replicate (Figure 2E). The relatively small variance of effect estimates, despite only being fitted with three replicates, and the strong posterior mean correlation between batches (Figure 2F, Data S2; *r >* 0.937*, p <* 5.99 · 10*^−^*^6^) confirm the reproducibility of villages and the power of our model to estimate donor proliferation. We also compared the per-donor posterior means estimated by Townlet to the arrayed exponential growth rates. There was high correlation (Figure 2G, *r >* 0.748*, p <* 0.007) even though arrayed estimates were from only two time points, while village estimates were from four.

We further evaluated Townlet’s ability to recover similar donor proliferation effects from repeat village experiments using predictive modeling. Using posterior predictive checks, thousands of new village experiments with the same design were simulated by sampling from the fitted model. The observed data was then compared to the simulated data to check if it fell into the bounds of the simulation. In both separately modeled batches, *>* 95% of the observed data fell within the 95% credible intervals of these new simulated distributions, thereby demonstrating accurate model estimation (Figure S1A-S1B). Additionally, we fit Batch 1 data and used it to predict the unobserved data from Batch 2 by simulating new villages from the fitted posteriors. Again, *>* 95% of the real experimental data from Batch 2 fell within these resulting posterior predictive distributions. These results demonstrate the importance of using a compositional model such as Townlet when assessing village proliferation dynamics.

### Townlet has enhanced power to detect factors that influence donor fitness

We performed a statistical power analysis to assess our Townlet’s ability to recover meaningful biological signals from *in silico* villages compared to off-the-shelf approaches (e.g., linear, frequentist Dirichlet, and exponential growth models; Methods ). Synthetic village data was generated using a simulation framework that we built using Townlet’s hierarchical Dirichlet regression model. Within this framework, we fixed each donor’s proliferation effects and generated new simulated village proportion data. We evaluated two model scenarios where we attempted to recover the simulated: (i) group proliferation effects from villages under control conditions and (ii) treatment-specific donor proliferation effects.

To address issue (i), we ran hundreds of *in silico* experiments where we varied the number of donors, replicates, and sample time points. When trying to recover donor proliferation group effects, Townlet out-performed all other methods–even when effects were small–and it adequately controlled false positive rate (Figure 3A-3B; Figure S2A-S2F). As expected, increasing the number of donors had the strongest effect on sensitivity (Figure 3A); however, increased experiment length (e.g., beyond 3 cell passages) did not increase sensitivity, likely due to faster donors dominating later sampled time points (Figure S2C).

**Figure 3.**
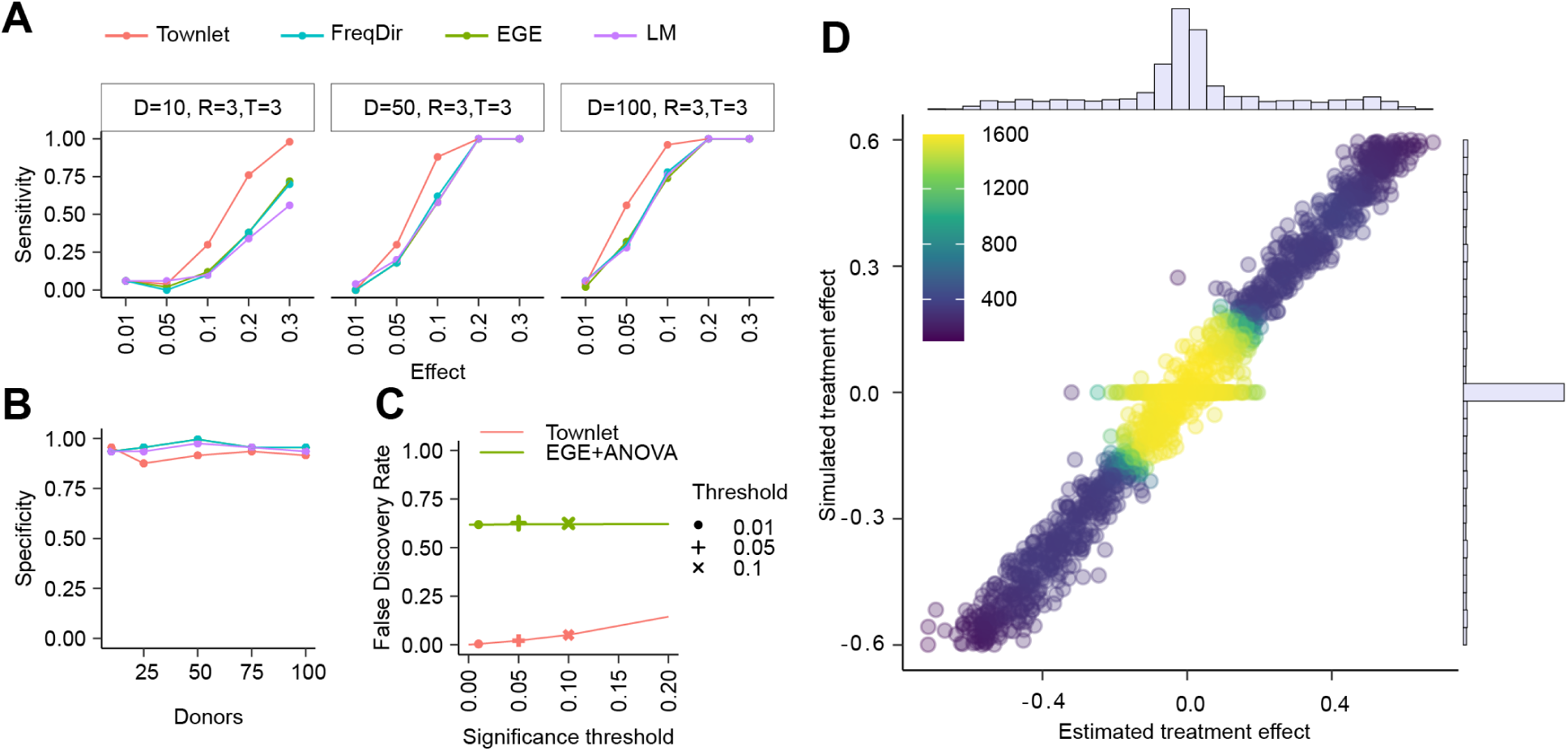
Townlet has higher power to detect proliferation effects compared to frequentist approaches. (A) Townlet has higher sensitivity, relative to other benchmarked models (freqdir = frequentist Dirichlet, exe = exponential growth equation, and lm = linear model), when performing inference on donor group effects (no treatment) using *in silico* village data (with R=3 replicates, T=3 sample time points) and varying donor numbers (D= 1, 50, 100). Increasing the number of donors in a village has the greatest influence on sensitivity. (B) Specificity is also high across simulated village data with varying numbers of donors. (C) Townlet has a much lower false discovery rate when detecting individual donor treatment effects compared to ANOVA based approaches. (D) Density plot showing correlation between simulated donor treatment effects and those estimated by Townlet. Note that 50% of the donors had their treatment effect simulated as zero and the remaining donor’s effects were drawn from a U(−0.6, 0.6).

Townlet also performed strongly when evaluating treatment-specific donor proliferation effects (issue (ii)). We simulated a generic treatment experiment where 3 simulated doses were applied to villages in triplicate and sampled over 3 time points. Our modeling approach better controlled false discoveries while maintaining high power and accuracy to detect effects (Figure 3C-3D). These simulations demonstrate the improved statistical power of Townlet’s modeling approach and the varied performance of statistical inference based on different village designs (e.g., number of donors, replicates, time points). With these results in mind, we applied the Townlet model on village-based fitness assays to map the genetics underlying NPC proliferation and vulnerability to environmental stress.

### Townlet reveals neurodevelopmental disorder-relevant mechanisms

We sought to determine if our village design and Townlet model could capture disorder-relevant group differences in proliferation. Cortical overgrowth and macrocephaly are commonly observed in individuals diagnosed with autism spectrum disorders (ASD), and there is evidence that hyperproliferative populations of fetal NPCs contribute to these features ^28–31^. For instance, individuals harboring the heterozygous deletion of the 16p11.2 chromosomal region (16pdel) are at higher risk of presenting with ASD, intellectual disability, and enlarged cortical surface area ^32–34^. Some studies have reported increased NPC proliferation in 16pdel models while others have not ^35–39^. The lack of consensus in human *in vitro* 16pdel studies is potentially due to heterogeneity across patient lines, small donor sample sizes (n = 3-4 patient lines), and the technical variation of arrayed approaches that can collectively mask the likely subtle, yet disorder-relevant differences in NPC proliferation rates between control and patient populations.

To reconcile these inconsistent observations, we constructed an NPC village consisting of 11 neurotypical controls and 12 patients that we sampled across 3 passages for Census-seq analysis (Figure 4A). Village well-to-well replicate correlation was high, and again, variance was lower in villages compared to array cultures (Figure S3A-S3B). Using Townlet, we calculated individual donor proliferation effects and village dispersion (Figure 4B; Figure S3C; Data S3). We detected a significant increase in 16pdel NPC proliferation relative to controls (*lfsr* = 4 · 10*^−^*^4^; Figure 4C). This result was consistent even when we excluded the fastest 16pdel donor (*lfsr* = 2 · 10*^−^*^4^, Figure S3D-S3E). Importantly, the individual donor proliferation effects estimated by Townlet correlated well with the individual proliferation rates estimated from arrayed cultures (Figure 4D, *r >* 0.761*, p <* 2.74 · 10*^−^*^5^). Further, deletion-control comparisons made from the arrayed data confirmed the significantly increased 16pdel NPC proliferation rates (*p* = 0.034, Figure 4E), as did an arrayed growth comparison between a control and 16p11.2 CRISPR-engineered isogenic line ^40^ (*p* = 8.21 · 10*^−^*^5^, Figure 4F). These results indicate that this rare copy number variant may contribute to macrocephaly through overgrowth of fetal NPC populations.

**Figure 4.**
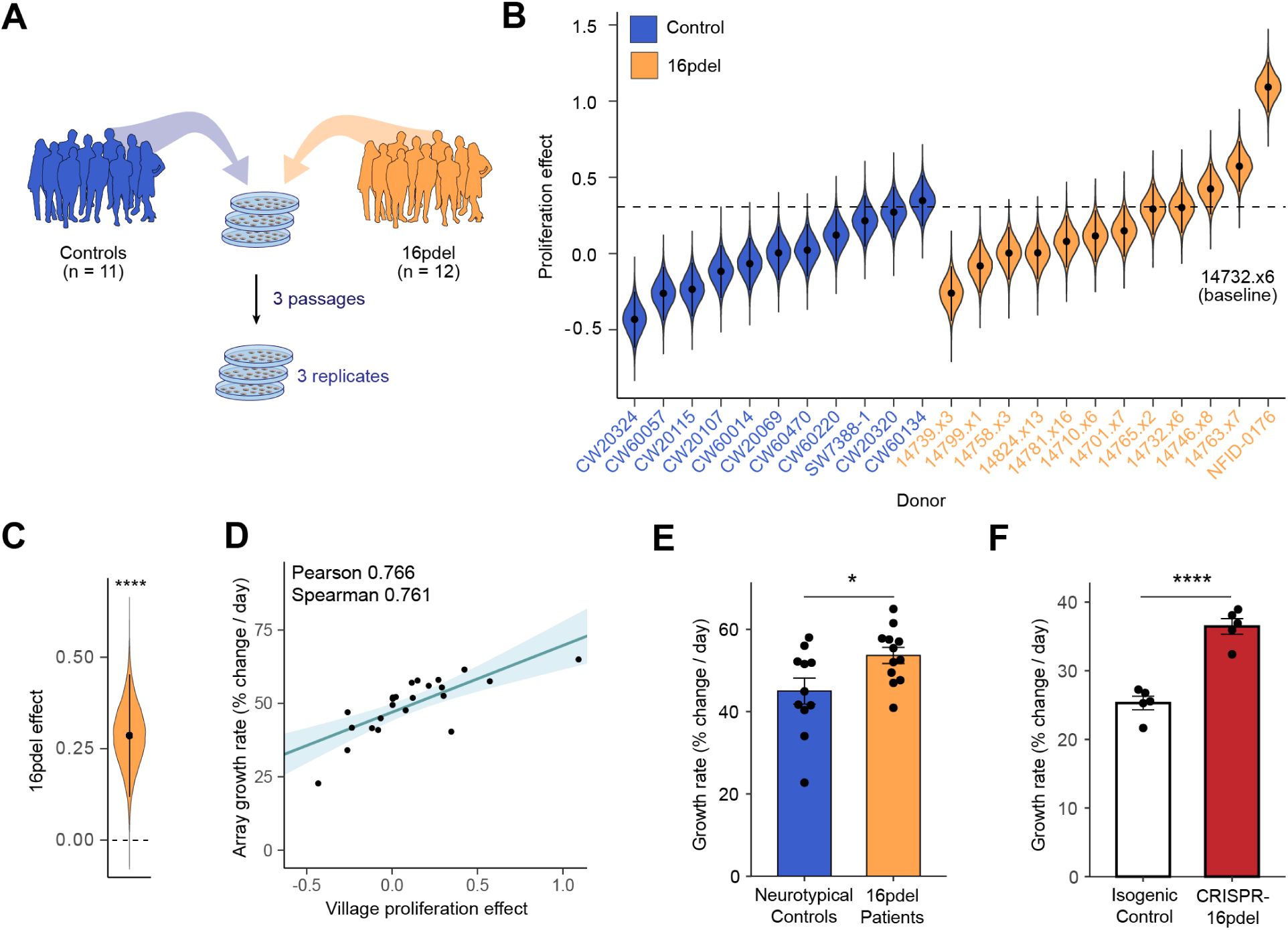
16p11.2 deletion NPCs are hyperproliferative. (A) Design of 16p11.2 deletion village. Twelve patient and eleven control NPC lines were pooled and plated in triplicate prior to harvest over three passages. (B) Total donor proliferation effects estimated from Townlet colored by donor deletion status relative to baseline donor 14732.x6 (represented as dashed line). All posteriors include the mean (point) and 95% credible interval error bars. (C) 16pdel patient group grew significantly faster than controls as shown by fitted posterior distribution being well above zero (*lfsr* = 4 *·* 10*^−^*^4^). (D) Proliferation effects from Townlet correlated strongly with exponential growth rates estimated from arrayed plates prior to pooling (n = 23 lines. Pearson r = 0.766, Spearman r = 0.761). (E) Growth rates estimated from arrays confirm 16pdel hyperproliferation. Mean +/- s.e.m. (t-test, p = 0.034). (F) CRISPR-engineered 16p11.2 deletion line grew significantly faster than isogenic control. Mean +/- s.e.m.(n = 5 wells per line; t-test, *p* = 8.21 *·* 10*^−^*^5^).

### Common genetic loci associated with NPC proliferation

Complex traits are shaped not only by rare, large-effect mutations like 16p11.2del, but also by common variants with small to modest impact. The genome-wide association study (GWAS) approach is routinely used to identify the genetic loci that have statistical relationships with human traits of interest, such as neuroanatomical features, behavioral tendencies, and risk for brain disorders. Human subject GWASs typically require thousands of participants due to the challenges with accurately quantifying traits and properly accounting for non-genetic factors and confounders (e.g., environmental exposures) ^41–43^. Remarkably, “GWAS-in-a-dish” experiments using arrayed culture designs have detected genome-wide significant loci for viral susceptibility ^15,44^, cell morphology ^45^, and drug response ^46^ from small sample sizes (∼40-300 donors). Here, we leveraged villages and the improved sensitivity and accuracy of Townlet to nominate genomic loci associated with between-donor line differences in NPC proliferation.

We constructed a village of 39 hPSC-derived NPCs, harvested samples in triplicate at five time points (Day 0, 1, 5, 6, and 10; passaged at Day 5; Figure 5A) for Census-seq, and generated a single proliferation measurement per donor using Townlet. We excluded one donor from analysis that showed lower than expected representation at Day 0 (*<* 0.5%). Further, we detected three sibling pairs in the village via whole genome sequencing, so we randomly removed one donor from each pair to minimize the effects of relatedness on GWAS analysis (Figure S4A). As expected, among the 35 remaining donors, we observed considerable variation across donors in proliferation (proliferation effect sizes = −0.41 to 0.56; Figure 5B).

**Figure 5.**
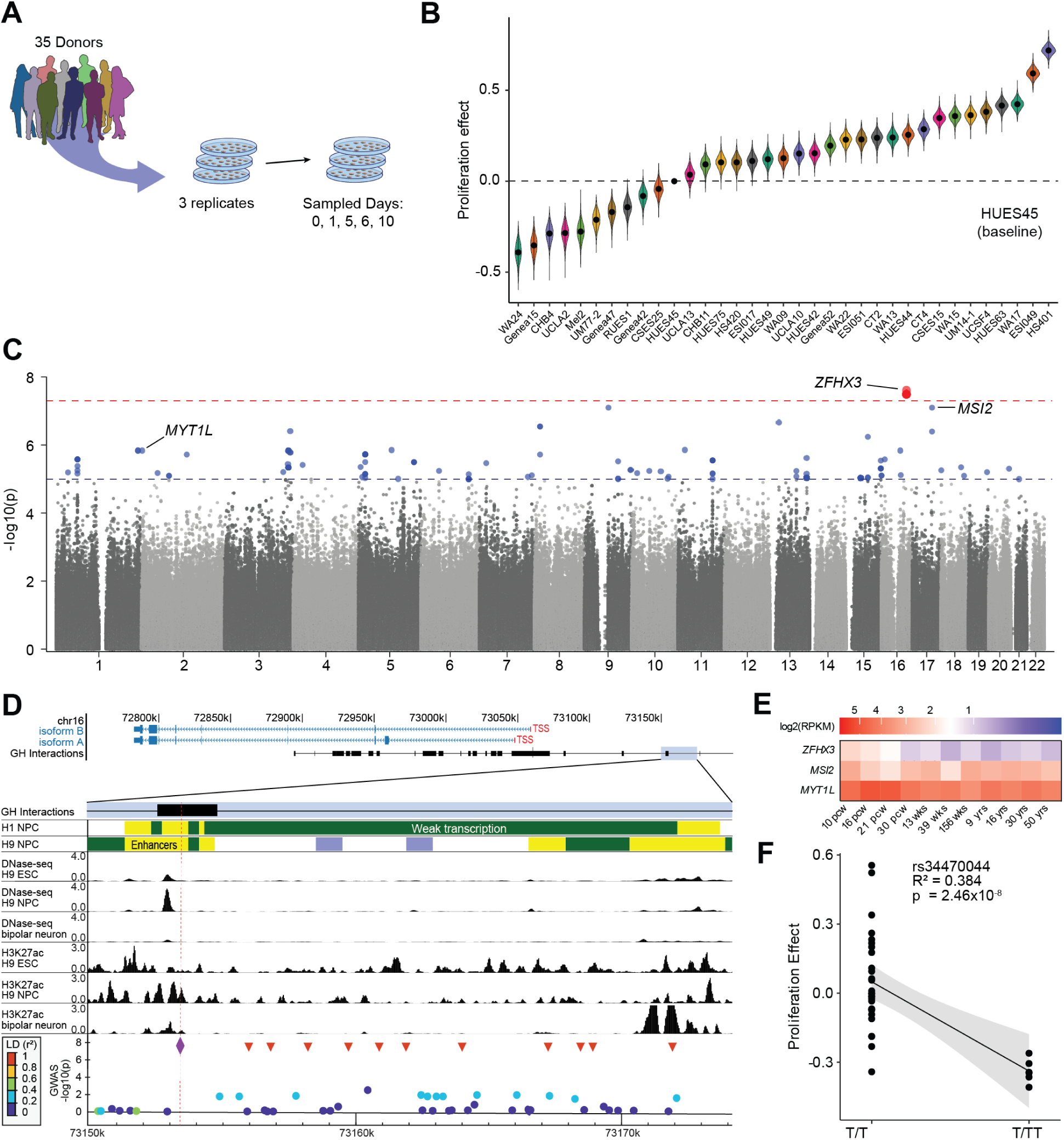
Common variants in *ZFHX3* gene are associated with NPC proliferation. (A) Thirty-nine NPC lines were pooled into a village, plated in triplicate, and sampled on Days 0, 1, 5, 6, and 10. Four donors were bioinformatically removed prior to downstream analysis. (B) Total proliferation effect posteriors estimated for the 35 donors grown in control conditions relative to the baseline donor HUES45 (points = means, error bar = 95% credible intervals). (C) Manhattan plot of genome-wide association scan for donor proliferation effects. Y-axis denotes – log_10_(p-value) from GEMMA likelihood ratio tests of SNP associations with the proliferation effect posteriors (biological n=35). Genome-wide significance threshold (*p <* 5 *·* 10*^−^*^8^) marked by red dashed line. Blue dashed line shows threshold for suggestive evidence (*p <* 1 *·* 10*^−^*^5^). (D) Regulatory context at the *ZFHX3* locus. (Top) Genomic locus with RefSeq isoform tracks ^56^. GeneHancer interactions (black box) denote high confidence enhancer-gene associations ^57^. (Bottom) Enlarged view of the blue-highlighted 25 kb region showing local epigenomic data. Chromatin state annotations from Roadmap Epigenomics Project ^58,59^ highlight predicted enhancers (yellow) and weak transcription regions (green) in hPSC-derived NPCs. DNase-seq and H3K27ac signal tracks indicate cell type-specific enhancer activity in NPCs ^60^. GWAS results were plotted using LocusZoom ^61^, with dot color indicating LD (r^2^) to the index SNP rs34470044 (chr16:73153445T:TT, *p* = 2.46 *·* 10*^−^*^8^; purple diamond, red dashed line). The locus contains 11 SNPs in near-perfect LD with the index SNP (red triangles). ESC = embryonic stem cells; GH = GeneHancer; TSS = transcription start site. (E) Developmental expression (log_2_ RPKM, color heat scale) for *ZFHX3*, *MSI2*, and *MYT1L* across 11 human brain stages ranging from 10 post-conception weeks to adulthood. Data from Brainspan ^62^. (F) Association of rs34470044 donor genotype with proliferation effect. The solid line shows the linear regression with a shaded band indicating the 95% confidence interval (R^2^ = 0.384).

Using Townlet proliferation effect sizes as the phenotype and whole genome sequencing data collected from each individual donor, we ran a GWAS using a linear mixed model to adjust for population structure and relatedness among individuals ^47^. Analysis of common variants (minor allele frequency *>* 5%) identified one locus (12 SNPs) that reached genome-wide significance in association with proliferation (*p <* 5 · 10*^−^*^8^) and nominated 50 loci (162 SNPs) with suggestive evidence (*p <* 1 · 10*^−^*^5^; Figure 5C; Data S4). Given the relatively small number of samples compared to large sample size human subject GWASs, we selected the top 100 loci ranked by p-value for downstream analysis. These loci predominantly mapped to non-coding regions of 49 protein-coding genes, many of which are expressed in the human fetal brain and *in vitro* NPCs ^15^ (Figure S4B). Notable genes include NPC proliferation modifiers *MYTL1*, *BBS2*, and *MSI2*, and gene ontology (GO) terms nervous system development (GO:0007399), regulation of stem cell proliferation (GO:0072091), and cell differentiation (GO:0030154) were significantly enriched after permutation testing (Figure S4C). These 49 genes were significantly enriched for association with neurodevelopmental disorder risk and tumor suppression ^48–55^(Figure S4D) supporting the clinical importance of dysregulated NPC proliferation. Our analyses confirm that these *in vitro* GWAS loci regulate genes with established roles in human prenatal neurodevelopment.

The genome-wide significant locus harbors 12 SNPs in Chromosome 16q22.3 that are in high linkage disequilibrium (LD; Figure 5D) and map to regulatory regions of the neurodevelopmental disorder gene *ZFHX3*, which is a transcription factor that regulates neuronal differentiation and acts as a cell cycle inhibitor ^63–66^). *ZFHX3* ChIP-seq peaks sit predominantly on promoters of neurodevelopmental genes in human NPCs ^63^) and expression is enriched in early prenatal stages in humans and other species when symmetrically dividing NPCs are abundant (Figure 5E). The lead SNP rs34470044 (chr16:73153445T:TT, *p* = 2.46 · 10^−8^) is in a region upstream of *ZFHX3* that is characterized by high chromatin accessibility and H3K27ac signals in hPSC-derived NPCs–but not hPSCs or post-mitotic neurons–and therefore shows hallmarks of a cell-type dependent enhancer (Figure 5D). Genotype at rs34470044 explains 38.4% of the inter-donor line variation in NPC proliferation with T/T donors showing increased proliferation relative to T/TT donors (*β* = −0.555, R^2^ = 0.384; Figure 5F). To validate this relationship, we calculated arrayed growth rates from an independent cohort of 114 NPC lines and confirmed T/T donors proliferated significantly faster than T/TT donors (*p* = 6.25 · 10^−3^; Figure S5E). Collectively, these results suggest that common genetic variants in *ZFHX3* regulatory regions may contribute to natural variation in cell-intrinsic human NPC proliferation rates.

### Between-donor differences in susceptibility to neurotoxicants

We further extended the utility of our village cultures and Townlet model to capture and explain treatment-dependent changes in cell viability across many donors. The World Health Organization ranks the heavy metal lead (Pb) as a top chemical of major public health concern due to its pervasiveness and effects on human health ^67,68^. The fetal brain is particularly sensitive to Pb, and early life exposure shows a continuous dose-response relationship with adverse neurobehavioral outcomes ^69^. Human subject genetic studies have nominated variants associated with elevated blood Pb levels ^70^) and risk of negative cognitive outcomes ^71^), but neither the extent to which individuals differ in Pb sensitivity at the cellular level nor the genetics underlying differential cell-intrinsic risk have been explored.

We first confirmed that our *in vitro* model is susceptible to Pb by exposing a single donor and observing dose-dependent reductions in viability using the CellTiterGlo2.0 reagent (Figure S5A). We then treated the 39-donor NPC village with 0, 3 *µ*M (high physiological dose), and 10 *µ*M (supraphysiological dose) Pb concentrations and sampled 4 and 7 days later using Census-seq (Figure 6A) ^72^. After excluding the same four donors from downstream analysis for low initial representation and relatedness, we used Townlet to integrate compositional data across all time points and concentrations into a single distribution of effect sizes that reflects Pb sensitivity for each donor relative to a baseline donor (CHB11, Figure 6B; Data S5). Donor Pb effect scores ranged from −0.54 (UM14-1) to 0.38 (ESI049), where more positive scores denote increased Pb resistance. Eleven donors were significantly more resistant than the baseline donor (*lfsr <* 0.05), and 11 donors were significantly more vulnerable (*lsfr <* 0.05). Dose-specific Pb effects calculated from Townlet showed similar between-donor variation (Figure S5B). An off-the-shelf, non-Dirichlet approach to calculate donor cell viabilities (Methods) confirmed the considerable between-donor variation in Pb response that ranged from 15.04% (donor: UM14-1) to 79.16% (donor: ESI049) viability one week after exposure (Figure S5C-S5D). These analyses reveal surprisingly high heterogeneity in cellular sensitivity to Pb across different genetic backgrounds and highlight the need to account for donor-intrinsic variability when performing toxicological risk assessments.

**Figure 6.**
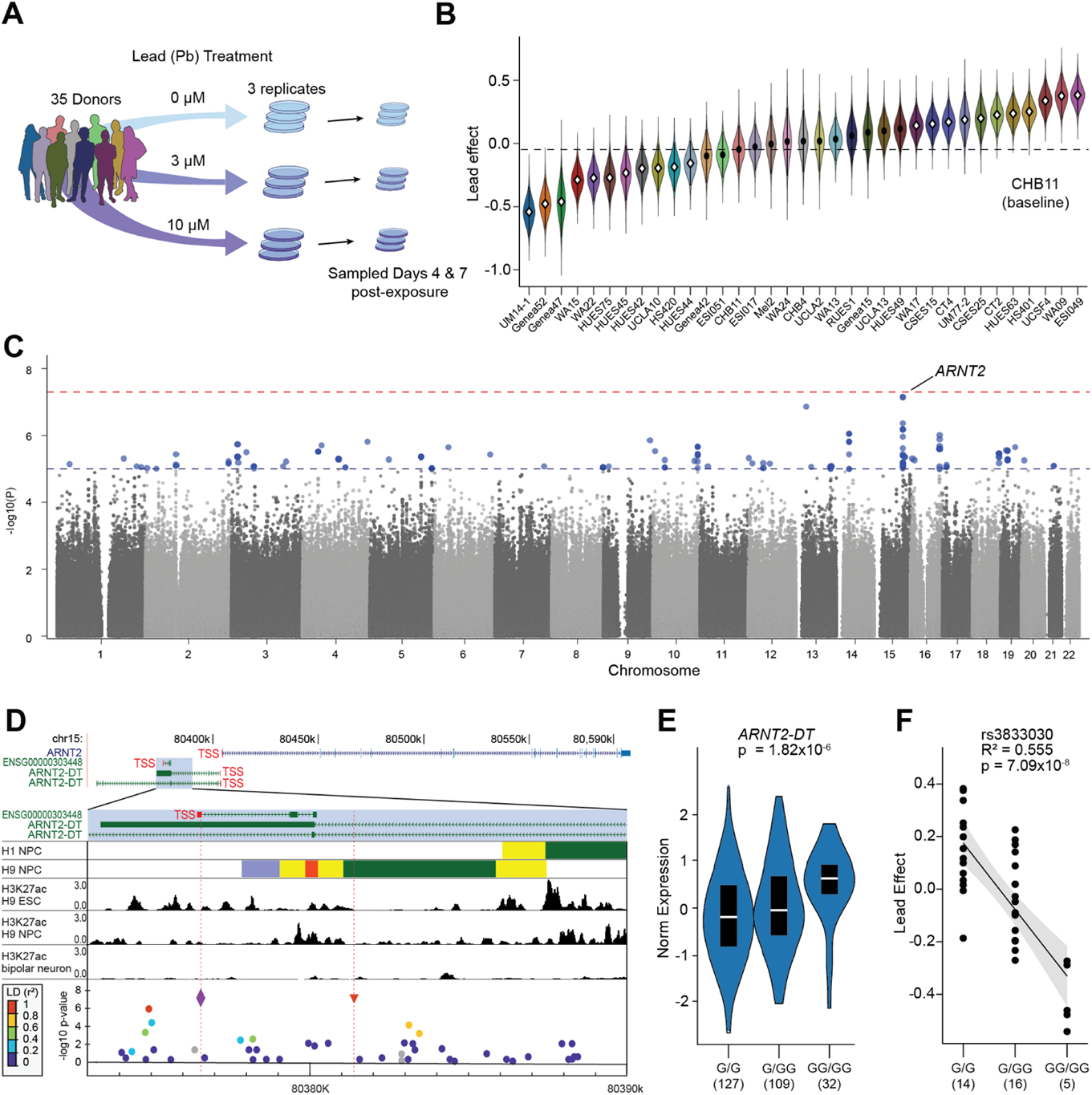
GWAS-in-a-dish analysis of lead (Pb) sensitivity. (A) Schematic of Pb exposure experiment. Thirty-nine NPC lines were pooled into a village and plated in triplicate for each dose of Pb treatment (0, 3, or 10 *µ*M). Cells were harvested on Days 4 and 7 post-exposure for Census-seq. Four donors were bioinformatically removed prior to downstream analysis. (B) Donor Pb effects estimated by Townlet across all treatment doses relative to baseline donor CHB11. Donors with white diamonds (posterior means) significantly differ from the baseline donor. More positive values denote increased resistance to Pb. (C) Manhattan plot. Y-axis denotes –log_10_(p-value) from GEMMA likelihood ratio tests of SNP associations with Pb effect posteriors (biological n = 35). Red line: Genome-wide significance (*p <* 5 *·* 10*^−^*^8^). Blue line: Suggestive significance (*p <* 1 *·* 10*^−^*^5^). The top association maps to a regulatory divergent transcript of *ARNT2*. (D) Regulatory context at the *ARNT2* locus. (Top) Genomic locus with RefSeq isoform tracks ^56^. (Bottom) Enlarged 17 kb region with local epigenomic data. The vertical red dashed lines mark the top SNPs rs3833030 (chr15:80376573G:GG, *p* = 7.09*·*10*^−^*^8^, purple diamond) and rs3858973 (*p* = 7.09*·*10*^−^*^8^, red triangle). rs3833030 resides 125 basepairs downstream of the transcription start site for a noncoding transcript, within the coding region of one *ARNT2-DT* isoform, and within the intron of another *ARNT2-DT* isoform. Roadmap Epigenomics chromatin state predictions are shown ^58,59^ as well as H3K27ac signal tracks from *in vitro* neural cell types ^60^. GWAS results were plotted using LocusZoom ^61^, with dot color indicating LD (r^2^) to rs3833030. (E) rs3833030 is an eQTL for *ARNT2-DT* in human frontal brain cortex. Violin plots show normalized expression grouped by genotype (*p* = 1.82 *·* 10*^−^*^6^, data from GTEX). Black boxes indicate expression quartiles and the inner white line marks the median expression per genotype. (F) Association of donor genotype at rs3833030 with Pb effects estimated by Townlet. Solid line shows linear regression with shaded 95% confidence interval (R^2^ = 0.555).

We performed a GWAS to identify common variants that explain between-donor variation in Pb effect scores. Though no loci reached the genome-wide significance threshold, there was suggestive evidence for 49 loci (334 SNPs; *p <* 1 · 10*^−^*^5^; Figure 6C). The top 100 loci mapped to 67 protein-coding genes that were enriched in such GO terms as cellular response to xenobiotic stimulus (GO:0051552) and AIM2 inflamma-some complex (GO:0140970; Figure S5E). This gene list also significantly overlapped with risk genes for neurodevelopmental disorders, implying that heavy metal exposure may disturb brain development through similar molecular pathways that underlie these conditions (Figure S5F).

The top locus includes 2 SNPs in perfect LD (rs3833030 and rs3858973, GWAS *p* = 7.09 · 10*^−^*^8^) located in a divergent transcript of the brain-enriched neuroprotective transcription factor *ARNT2* (Figure 6D-6E). In our data, genotype at rs3833030 explains 55.5% of the between-donor line variation in Pb vulnerability (*β*=-0.258, R^2^=0.555; Figure 6F). This response was dependent on risk allele copy number: donors with zero copies (G/G) were most resistant, followed by heterozygotes (G/GG), and then homozygous carriers (GG/GG). These results indicate that natural variation in *ARNT2* -mediated processes may contribute to donor differences in Pb-induced neurotoxicity.

## Discussion

*In vitro* fitness assays are widely used to assess how cells proliferate, survive, and respond to changing environments. Most studies are performed on a few donor lines representing limited genetic variation due to the limitations of conventional methods, many of which we overcome by taking a village-based approach. The co-culture of donors, however, introduces certain considerations that we address in this work. First, a common concern is the potential influence of non-self on *in vitro* donor line traits. We show that villages successfully capture donor-intrinsic proliferation rates in a reproducible manner, indicating that non-cell autonomous influences are minimal. These findings support work that found strong similarities between array- and village-based assays of viral infection susceptibility ^15^ and gene expression ^16^. Second, we present a Bayesian Dirichlet model named Townlet for time-series cell fitness analysis that models the between-donor dependencies expected in Census-seq data. To address the limited number of replicates common in genomic experiments, Townlet integrates information via multi-dimensional shrinkage across donors, time points, and treatments to provide robust, low-variance estimates.

With these tools, we performed case-control comparisons to understand the connection between 16p11.2 deletion and macrocephaly. Recent human *in vitro* studies have suggested overgrowth of NPC populations associated with 16pdel; one group reported patient stem cell-derived NPC hyperproliferation (n = 2 donors) using cell counting and EdU incorporation assays ^38^, while another found enlarged neural rosettes in ventral organoids generated from a CRISPR-engineered isogenic stem cell line ^39^. Other studies found no differences in NPC cell cycle dynamics, and instead nominated enlarged soma size, increased dendritic area, and reduced phagocytosis as contributing factors to brain overgrowth in patients ^35,36,38,73,74^. Our village analysis–which represents the largest 16pdel *in vitro* study to date in terms of donor sample size–indicates that overgrowth of early fetal NPC pools contributes to macrocephaly. Previous studies have reported NPC hyperproliferation in human *in vitro* ASD models ^23,75,76^, suggesting this may be a node of biological convergence across genetic risk factors. Our results and others ^77^ highlight the utility of villages for future case-control comparisons.

Our focus on one cell type combined with low technical noise and a refined computational model likely contributed to our ability to detect what is, to our knowledge, the first *de novo* genome-wide significant association discovered using cell villages. The 35-donor GWAS for proliferation produced a genome-wide significant locus (index SNP: rs34470044) that maps to a distal enhancer of *ZFHX3*. In the rodent brain, *ZFHX3* overexpression results in NPC cell cycle arrest, while low protein abundance in the nucleus is characteristic of proliferating NPCs in the ventricular zone ^64^. Our outcomes suggest *ZFHX3* influences naturally occurring variation in proliferation dynamics across human donors, which could have wider implications on diversity in brain traits like cortical surface area and cognition ^78^. Future studies elucidating the exact mechanisms underlying this genotype-phenotype relationship are warranted.

We also nominate genetic modifiers of cell-intrinsic vulnerabilities to environmental exposures, which is an area where human subject GWASs have largely fallen short due to limited cellular resolution and reliance on imprecise exposure estimates (e.g., zip code-level data) ^43,79^. In our study, we identified a locus upstream of *ARNT2* that was just below the genome-wide significance threshold (index SNP: rs3833030). ARNT2 forms heterodimers with bHLH-PAS sensor proteins like HIF-1*α* to regulate gene expression in response to environmental stimuli such as oxidative stress and hypoxia ^80^. Though its role in Pb response has not been studied, it is possible that ARNT2 protects cells from the pseudo-hypoxic conditions and production of reactive oxygen species observed in Pb-exposed neural cells ^81^. Our work introduces ARNT2 as a potential therapeutic target for mitigating heavy metal cytotoxicity and paves the way for future village-based efforts to dissect cell-intrinsic vulnerabilities to the thousands of chemicals that comprise the human exposome ^82,83^. Our study aims to push the boundaries of village experiments and empower future efforts to explain how human genetic variation confers phenotypic diversity in both healthy and disease states. Several additional improvements to our system could facilitate this vision. First, use of cell culture robotics could increase the number of donors and ancestries represented in a village, which would increase statistical power. Second, as villages scale to hundreds of donors, approaches such as combinatorial pooling or meta-analysis of multiple villages will be critical to maintain power and reproducibility, and should be systematically evaluated through both experimentation and simulation ^19^. Third, more comprehensive simulation frameworks, akin to those used in single-cell transcriptomics ^84^, could help optimize experimental design by clarifying how parameters such as donor diversity, culture duration, and environmental perturbations influence sensitivity. Finally, extensions of Townlet that enable proper analysis of FACS-based phenotypes, as we recently reported using off-the-shelf tools ^15^, could increase the number of phenotypes that can be assayed in a village and expand opportunities for discovery.

## Resource Availability

### Data availability

Users can apply for controlled access to genotype data (VCF files) for the hESCs donors (e.g., Proliferation and lead GWAS villages) at https://duos.broadinstitute.org (Accession number DUOS-000121). Once approved, the data can be downloaded from this Terra workspace: https://app.terra.bio/#workspaces/convergneuro-mccarroll-anvil/Broad ConvergentNeuro McCarroll Nehme hESC HMB VillageData. Raw low-coverage whole genome sequencing data used for Census-seq input can be accessed at the database of Genotypes and Phenotypes (dbGap) using accession number [IN PROGRESS]. The Census-seq readouts used as input to the Townlet model can be found in the Supplementary Data Files.

### Code availability

Townlet is available to run as an R package and can be installed from https://github.com/pimentellab/townlet. Code used for simulations and Townlet analyses can be found at https://github.com/hansonch/townlet analysis.

## Acknowledgements

This work was supported by NIH grant R00MH119327 (M.F.W.), HHMI Hanna H. Gray Fellowship (H.P.), Hypothesis Fund (H.P.), the UCLA Eli and Edythe Broad Center of Regenerative Medicine and Stem Cell Research Award (P.A., M.F.W.), the California Institute for Regenerative Medicine DISC0-15737 (M.F.W.), the CIRM UCLA Eli and Edythe Broad Center of Regenerative Medicine and Stem Cell Research Training Program (R.G.F.), the Dana Foundation (A.R.V.), and the NHGRI Genomic Analysis and Interpretation Training Program (L.S.). C.H. is supported by the NIH National Human Genome Research Institute (NHGRI) grant F31HG013890. The content is solely the responsibility of the authors and does not necessarily represent the official views of the NIH. We thank Drs. Michael Talkowski (Masschusetts General Hospital) and Xander Nuttle (University of Michigan) for providing the isogenic 16pdel lines, and Bogdan Pasaniuc (University of Pennsylvania) for constructive feedback on this manuscript.

## Author contributions

Conceptualization and design, C.H., T.D., P.A., H.P., and M.F.W.; experimentation, T.D., A.R.V., Y.S.K., H.L., L.S., T.E.D.; Townlet model, C.H. and H.P.; data analysis, C.H. and T.D.; writing, C.H., T.D., A.R.V., Y.K.S., H.P., and M.F.W.; supervision, H.P.. and M.F.W.

## Declaration of generative AI and AI-assisted technologies

During the preparation of this work, the authors used ChatGPT from OpenAI to consolidate text. After using this tool, the authors reviewed and edited the content as needed and take full responsibility for the content of the published article.

## Supplementary Information

Document S1. Figures S1–S5 Table S1: Full list of human pluripotent stem cell lines used for each experiment.

Table S2: Meta-data for human pluripotent stem cell lines.

Data S1: Census-seq outputs for village-based fitness assay validation.

Data S2: Townlet proliferation effect scores for village-based fitness assay validation.

Data S3: 16p11.2del village Census-seq and Townlet output.

Data S4: Proliferation GWAS.

Data S5: Lead GWAS.

## Methods

### Stem cell lines and maintenance

All protocols were approved by the UCLA Institutional Biosafety Committee (IBC) and the UCLA Human Pluripotent Stem Cell Research Oversight (hPSCRO) Committee. All hPSC methods were carried out in accordance with UCLA IBC and hPSCRO guidelines and regulations. Informed consent was obtained for all cell lines by the labs and/or biobanks that supplied the lines to the Wells lab. All human embryonic stem cells (ESCs) used in this paper are registered in the NIH Human Embryonic Stem Cell Registry. All neurotypical control human induced pluripotent stem cell (iPSC) lines were obtained from CIRM/FujiFilm Cellular Dynamics, except for SW7388-1, which was reprogrammed at the Broad Institute by Dr. Ralda Nehme. All 16p11.2 deletion patient iPSC lines were obtained from the Simons Foundation Autism Research Initiative (SFARI), except for NFID 0176, which was reprogrammed at the Broad Institute by Dr. Ralda Nehme. Isogenic unedited and 16p11.2 deletion lines were generously supplied by Dr. Michael Talkowski (Massachusetts General Hospital). Prior to receipt, all lines were confirmed via genotyping, karyotyping, and *in vitro* differentiation to establish pluripotency. The list of donor lines used for each experiment can be found in Table S1.

All hPSCs were maintained on Geltrex basement membrane matrix (1:100; Life Technologies, A1413301) and fed daily with mTeSR Plus media (Stem Cell Technologies, 100-0276). Upon reaching 80-90% confluency, cells were split by incubating for 10 minutes at 37°C in Accutase (Innovative Cell Technologies, AT104) followed by a 1:1 dilution in mTeSR Plus and centrifugation. Cells were resuspended in mTesR Plus with ROCK inhibitor Y-27632 (10 *µ*M; Stemgent, 04-0012) and plated at 1:10 to 1:30 dilution. ROCK inhibitor was removed after 24 hours.

### Stem cell-derived neural progenitor cell (NPC) induction and maintenance

Human PSCs were transduced and induced into NPCs as previously described using the SNaP method ^15,26^. In brief, human stem cells were transduced with TetO-Ngn2-Puromycin and Ubq-rtTA lentiviral constructs. For SNaP generation, transduced hPSCs were plated on Geltrex in mTeSR Plus supplemented with ROCK inhibitor Y-27632 (10 *µ*M; Stemgent, 04-0012). The next day, cells were fed with Induction Media (Day 1): DMEM/F12 (ThermoFisher, 11320082), Glutamax (1:100; ThermoFisher, 10565018), 20% Glucose (1.5% v/v), N2 Supplement (1:100, ThermoFisher, 17502048), Doxycycline (2 *µ*g/mL; Sigma-Aldrich, D9891), LDN-193189 (200 nM; Stemgent, 04-0074), SB431542 (10 *µ*M; Tocris, 1614), and XAV939 (2 *µ*M; Stemgent, 04-00046). Twenty-four hours later, cells were fed with Selection Media (Day 2): DMEM/F12, Glutamax (1:100), 20% Glucose (1.5% v/v), N2 Supplement (1:100), Doxycycline (2 *µ*g/mL), puromycin (5 *µ*g/mL; ThermoFisher, A1113803), LDN-193189 (100 nM), SB431542 (5 *µ*M), and XAV939 (1 *µ*M). After 24 hours in Selection Media, cells were replated in NPC maintenance media supplemented with puromycin (5 *µ*g/mL) and Y-27632 (10 *µ*M). Starting 12–18 hours post-passaging, cells were maintained in NPC maintenance media without puromycin or Y-27632. Immunostaining was performed to validate NPC identity, defined as ≥75% NESTIN+/PAX6+/SOX1+ and ≤ 0.1% OCT4+ cells at Passage 1-3.

NPCs were maintained on Geltrex basement membrane matrix (1:100; Life Technologies, A1413301) and fed daily with NPC maintenance media, which was made fresh every 48 hours: DMEM/F12 (Thermo Fisher Scientific; 11320082), Glutamax (1:50; Thermo Fisher Scientific, 10565018), MEM-NEAA (1:100; Life Technologies, 10370088), Pen/Strep (1:100, Cytiva, SV30010), B27 minus Vitamin A (1:50; Life Technolo-gies, 12587010), N2 Supplement (1:100; Life Technologies, 17502048), recombinant human EGF (10 ng/mL; STEMCELL Technologies, 78006.1), recombinant human basic FGF (10 ng/mL; Thermo Fisher Scientific, PHG0368). Cells were split weekly by incubating for 10 minutes at 37°C in Accutase (Innovative Cell Tech-nologies, AT104) and plating at 120,000 cells/cm^2^. For the first 24 hours after every passage, media was supplemented with ROCK inhibitor Y-27632 (10 *µ*M; Stemgent, 04-0012). After induction and expansion, NPCs were banked in Cryostor-10 (Stem Cell Technologies, 07930) for future use. Passage 2-8 NPCs were used for experimentation.

### Village Construction

NPCs from multiple donors were thawed, plated, and maintained as independent arrayed cultures for 5-7 days until confluent. Cell lines were frequently imaged to confirm healthy NPC morphology and were passaged at least once prior to village construction. To build villages, cells from each donor were dissociated with Accutase, resuspended in NPC maintenance media with ROCK inhibitor Y-27632 (10 *µ*M), and counted using a Countess II FL (ThermoFisher Scientific). Donors were pooled at equal proportions in a 50 mL falcon tube and then plated into replicate wells in 6-well plates at 40,000 cells/cm^2^ (with the exception of the Village versus Array experiment; see below).

### Census-seq

On harvest days, 250,000-500,000 cells per sample were pelleted and frozen at −80°C. DNA was then extracted from each sample following the standard procedure of the DNeasy Blood & Tissue Kit (Qiagen, 69504). All extracted DNA samples were indexed during library preparation, pooled, and sequenced using an Illumina Novaseq 6000 SP flow cell (paired end 150 bp reads; ∼20 million paired reads per sample).

### Village versus array comparisons

#### Village

We pooled 12 human ESC-derived NPC lines into a village at equal proportions and plated into six techni-cal replicate wells at 22,500 cells/cm^2^. The next day, cells from three of these wells were harvested for DNA extraction and Census-seq analysis in triplicate. The remaining three village wells were independently pas-saged every 5 days for a duration of 15 days. The passage 1 initial (day 1) and final (day 5) total cell counts were measured with a Countess Automated Cell Counter. These total cell counts were then multiplied by Census-seq’s corresponding donor composition estimates to calculate the cells per donor.

#### Array

Immediately following village construction, the remaining cell suspensions were independently passaged into 20 wells per donor in 96-well plates at a density of 3,333 cells/cm^2^. Half of the wells were stained with Hoechst-33342 (5 *µ*g/mL) and imaged using a BioTek Cytation 5 (Agilent Technologies; 10x Objective, 3×3 grid) on Day 1; on Day 5, the remaining wells were stained and imaged using the same procedure. CellProfiler ^85^ was used to quantify Hoechst-positive cells for both time points.

#### Exponential growth rates

For each array and village donor replicate, the exponential growth equation,

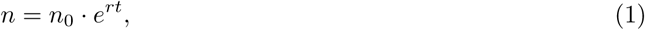

was used to estimate the percent change per day,

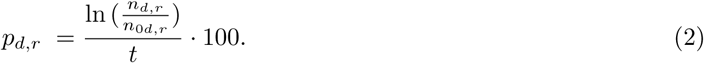

The final donor (d) cell count per replicate (r) is represented by *n_d,r_*, the initial donor cell count (*n*_0_*_d,r_*), and *t* is the time that elapsed between cell count measurements. The calculated *p_d,r_* array and village replicate values were then averaged.

### Hierarchical Dirichlet regression donor proliferation model

Townlet models donor proportion data using the Dirichlet distribution, which models a multivariate outcome that sum to one. The following parameterization was implemented:

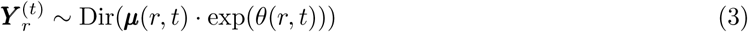

The ***Y*** ^(*t*)^ vector, of length D (number of donors), is our response variable that contains the compositional data of each donor (d) included in the village estimated from the Census-seq pipeline for a given time point (t) and replicate (r).

#### Estimating biological parameters

The first part of Equation 3 helps us characterize the biological dynamics within the experiment with

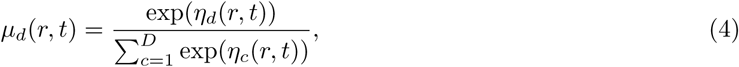

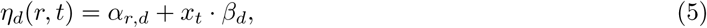

with priors,

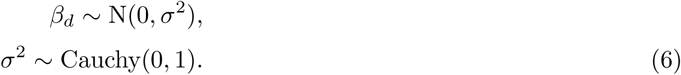

Here, each donor’s replicate specific proportion is modeled by *µ_d_*(*r, t*) using a softmax transformation to normalize all model components to unity. The donor specific model in log-space uses a log-linear model on *ν_d_*(*r, t*) which has an initial representation, *α_r,d_*, that is directly given to the model and a temporal proliferation effect, *β_d_*, that is estimated using the time index for each sample provided (*X_t_*, days from t0 or passage number).

The proliferation model can optionally incorporate treatments (e.g., chemical, pharmaceutical, environ-mental, etc.) and donor group covariates with interaction terms (e.g., sex, ancestry, phenotypes, disease status, sex:phenotype, sex:treatment, etc.) to infer significant influences on donor proliferation dynamics.

We can extend the model parameterization to include these effects

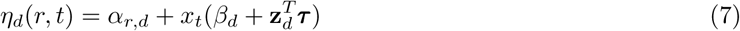

where the treatment effects (*τ_d_*) are partially pooled by donor,

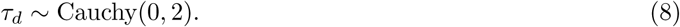

and the donor group effects (*τ_g_*) are pooled using the following prior,

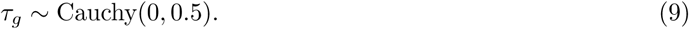

The Cauchy distributions provide shrinkage of effects towards zero, while still preserving large effect sizes with the heavy tails ^86^. The design matrix (*z_d_*) includes donor group identity and treatment dose information per donor. When donor group or treatment effects are included the total donor proliferation effect can be interpreted as *β_d_* + **z***^T^ **τ*** . All donors are relative to a baseline donor with a median proliferation rate since the prior on *β_d_* is centered at zero.

#### Accounting for overdispersion

The second component of equation 3 (*θ*(*r, t*)) accounts for overdispersion in individual village replicates over time. The following overdispersion parameters quantify technical variation that occurs during cell village experiments (e.g., uneven handling of samples during cell culture, sequencing bias, etc.). It is expected that the effects of technical variation will accumulate over time and may differ slightly across replicates, therefore, overdispersion is modeled by

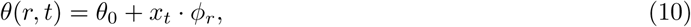

where,

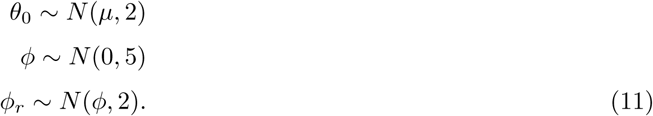

The amount of technical variation at village initiation is defined as *θ*_0_ and *ϕ_r_* defines the rate of change in variation over time per replicate. The intercept prior mean (*µ*) is estimated using maximum likelihood under the frequentist model.

### Simulate new cell village experimental designs using our generative Dirichlet model

*In silico* village composition data can be generated by: (i) fitting experimental data using our Dirichlet regression model and (ii) scaling and sampling from the interpretable fitted parameter posteriors to simulate new data. The following model parameters can be adjusted to simulate different village designs:

- **Donor proliferation effects (***β_d_, τ_d_, τ_g_*): The posteriors can be sampled and scaled to create infinite numbers of new *in silico* donors with optional treatment and donor group effects.
- **Initial village composition(***α_r,d_*): Randomly sample starting donor composition per replicate.
- **Sample time points and replicates (***x_t_*): The index can be modified to include more replicates and sample time points.
- **Dispersion parameters (***θ*_0_, *ϕ_r_*): Sample from and scale dispersion parameter posteriors to simulate villages with differing amounts of technical variation.

After setting, sampling and scaling all necessary parameters the mathematical framework from our Dirichlet regression model can be used to simulate new data. First, the *α* values for each donor are cal-culated and then these concentration parameters are used to randomly generate new representation data from a Dirichlet distribution. Repeating the simulation over multiple iterations (i.e., thousands of *in silico* experiments) gives us distributions of possible donor representation values under repeated experiments.

### Benchmarking Regression Models

Using *in silico* data from our statistical framework, we benchmarked Townlet against more naive analysis approaches. First, we tested the performance of our Dirichlet regression model when only a single donor covariate was included and no treatments. We benchmarked this version of the model against: (i) the exponential growth equation (EGE), (ii) a linear model (LM) and (iii) frequentist Dirichlet regression. These methods were implemented to estimate individual donor growth rates and then t-tests were performed to check for significant donor covariate effects. For Townlet, we used the local false sign rate ^87^ and the estimated effect direction of the posterior to assess if we could recover the simulated donor group effect. Sensitivity and specificity were calculated to compare performance between all four models.

Second, we benchmarked Townlet’s performance when estimating treatment effects per donor (no donor covariates). We compared our model to a more naive analysis approach where first EGE proliferation rates were individually estimated per donor. Then ANOVA followed by estimated marginal means testing was performed to identify donors whose average treatment growth rates significantly differed from the mean across all donors. This is analogous to our treatment effect (*τ_d_*). When simulating the data we set 50% of the donors to have a zero treatment effect and the remaining had a treatment effect drawn from

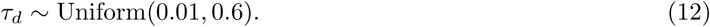

After running Townlet and this other approach on the simulated data, we used the false discovery rate to assess performance.

### 16p11.2#deletion proliferation

We constructed a 23-donor human iPSC-derived NPC village composed of 11 controls and 12 16p11.2 deletion donors. NPCs were thawed and maintained in three wells each of a 12-well plate before pooling, plating in triplicate at 40k/cm^2^, and passaging every 5 days for a total of 15 days. For each sample, at least 500,000 cells were pelleted prior to DNA extraction for Census-seq.

To validate that intrinsic NPC growth rates for 16p11.2 donors and neurotypical controls were retained in the village, the growth rates of the 23 donors were estimated from arrayed maintenance wells prior to pooling. Brightfield images (10x Objective, 3×3 grid) of these arrayed wells were captured using a Cytation 5 imager on days one and four post-plating (68 hours apart). Per well cell counts at each time point were then measured from these images using Cellpose ^88^. Exponential growth curves were independently fit for each well across all time points.

CRISPR-engineered isogenic NPCs carrying a 16p11.2 deletion and matched controls were plated at 40,000 cells/cm^2^ in five replicate wells each. Brightfield images (10x Objective, 3×3 grid; Cytation 5) were collected daily for six days. Cell counts were extracted using Cellpose, and exponential growth curves were independently fit for each well across all time points.

### Design of 35-donor NPC village

We pooled 39 human ESC-derived NPC donor lines into a village at equal proportions and plated in triplicate at 40,000 cells/cm^2^. Replicate wells were independently maintained and harvested for DNA sequencing at five time points (Day 0, 1, 5, 6, and 10; passaged at Day 5). For quality control, we applied the “Roll Call” algorithm from the Census-seq toolkit to estimate donor abundance at Day 0. One donor was excluded from subsequent analyses due to underrepresentation at this time point (*<* 0.5%). To limit potential confounding from sibling relatedness in downstream GWAS, we randomly excluded one donor from each of the three sibling pairs before estimating proliferation effects with Townlet and performing GWAS.

### GWAS analysis

We conducted genome-wide association analyses using Townlet outputs as the phenotypes and 30X whole-genome sequence (WGS) data from all donor lines in the village. Analyses were carried out using BCFtools (v1.11), PLINK (v2.0.0), and GEMMA (v0.98.5). We created a variant calling format (VCF) file that included the 35 donors and converted to PLINK binary format. Autosomal variants were retained, and SNPs were filtered to exclude markers with minor allele frequency (MAF) *<* 5% or genotype missingness *>* 5%. To account for population structure and relatedness, we generated a linkage disequilibrium (LD) pruned set of autosomal variants (window size = 50 SNPs, step size = 5 SNPs, r^2^ threshold = 0.2). LD was computed from the European population panel of the 1000 Genomes Project.This pruned dataset was used to compute the genetic relatedness matrix (GRM) in GEMMA. We performed association analysis across the full set of autosomal SNPs using GEMMA (linear mixed model, option −lmm 4), providing the proliferation effects estimated by Townlet as the primary phenotype alongside GRM and sex as covariates. After association testing, we performed clumping to identify approximately independent signals. Association summary statistics were processed in PLINK using an r^2^ threshold of 0.1 and a 250 kb window.

To prioritize candidate genes, we extracted the top 100 clumped variants ranked by likelihood ratio test p-value. These were mapped to gene symbols using the Ensembl Variant Effect Predictor. We then conducted Gene Ontology (GO) term enrichment via the gprofiler2 R package. Multiple testing correction was applied using the Benjamini–Hochberg false discovery rate method. For GO term permutation testing, we repeated the entire association workflow more than 500 times, each time randomly reassigning the Townlet-estimated proliferation effects across donors while keeping the covariates fixed.

Isoform and transcript annotations were obtained from the UCSC Genome Browser ^89^ using NCBI RefSeq gene models ^56^. Predicted enhancer–gene interactions were taken from GeneHancer ^57^. Chromatin state annotations were obtained from the Roadmap Epigenomics Project 15-core model ^57^. DNase-seq and H3K27ac ChIP-seq signal p-value tracks from human ESCs, NPCs, and neurons were downloaded from ENCODE ^60^ and visualized using the WashU Epigenome Browser ^90^. Association statistics were visualized with LocusZoom ^61^.

### Lead (Pb) treatment

To prepare a 50 mM lead (Pb) stock solution, 190 mg of lead(II) acetate trihydrate (Sigma, 316512) was dissolved in 10 mL of double-distilled water and subsequently diluted to the desired concentrations using NPC maintenance media. For array culture assays, NPCs derived from the Genea52 ESC line were plated at a density of 60,000 cells/cm^2^ into a 96-well plate. Five different Pb concentrations were tested (0, 1, 5, 10, 30 *µ*M), each with three replicates. Beginning the day after plating, culture media was replaced daily and supplemented with Pb. After 7 days of continuous exposure, cell viability was measured via CellTiter-Glo^®^ 2.0 reagent (Promega, G9241). Luminescence (ATP) readings were recorded for each well using a Cytation 5 imager and normalized to the mean luminescence of the 0 *µ*M (untreated) control replicates to determine relative viability.

For village experiments, replicate wells were treated daily with one of three Pb concentrations: 0, 3, and 10 *µ*M. Cells were harvested for DNA extraction and Census-seq in triplicate on Days 4 and 7 post-exposure. The samples underwent 0.5X low-coverage sequencing through Gencove (Long Island City, NY). Libraries were prepared using the Illumina Nextera DNA Flex Library Preparation Kit, then quantified by fluorometry. Quality assessment of the pooled libraries was performed using an Agilent TapeStation. Standard Illumina adapter sequences were employed and sequencing took place on an Illumina NovaSeq 6000 (paired end 150 bp reads: ∼6 million paired reads per sample).

To calculate donor response as percent viabilities, we first estimated the total number of cells in each replicate by using Cellpose analysis of brightfield images taken with a Cytation 5 imager (10x objective, 5×5 grid). Then, for each donor, we calculated an inferred cell count by multiplying its Census-seq derived proportion by the total cell count in that replicate. Finally, we determined each donor’s viability in a given replicate by dividing its inferred cell count in that replicate by its average cell count in the three untreated control replicates.

