## Supplementary Figures for "Cell villages and Dirichlet modeling map human cell fitness genetics"

### Supplemental Figures

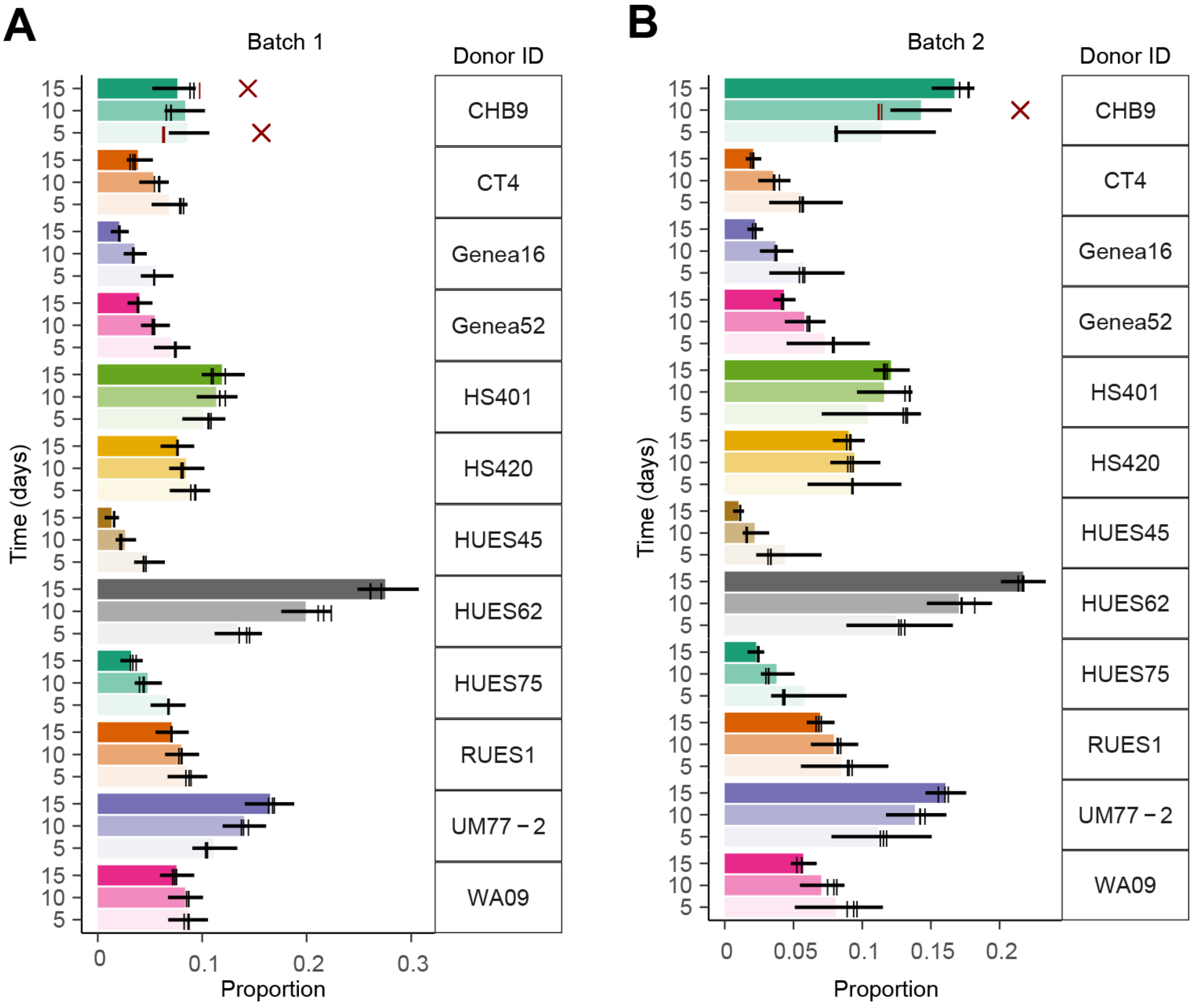


**Figure S1 Townlet posterior predictive checks**
(A-B) Posterior predictive checks simulate a distribution of representation values per donor and sample using the model fit (bars represent distribution mean with 95% credible interval errorbars). We were able to recover > 95% of the training data (tick marks, black) within these distributions in a, Batch 1 and b, Batch 2. Red Xs denote donors where at least one of the training data points did not fit (tick mark, red) within the posterior predictive check credible intervals.

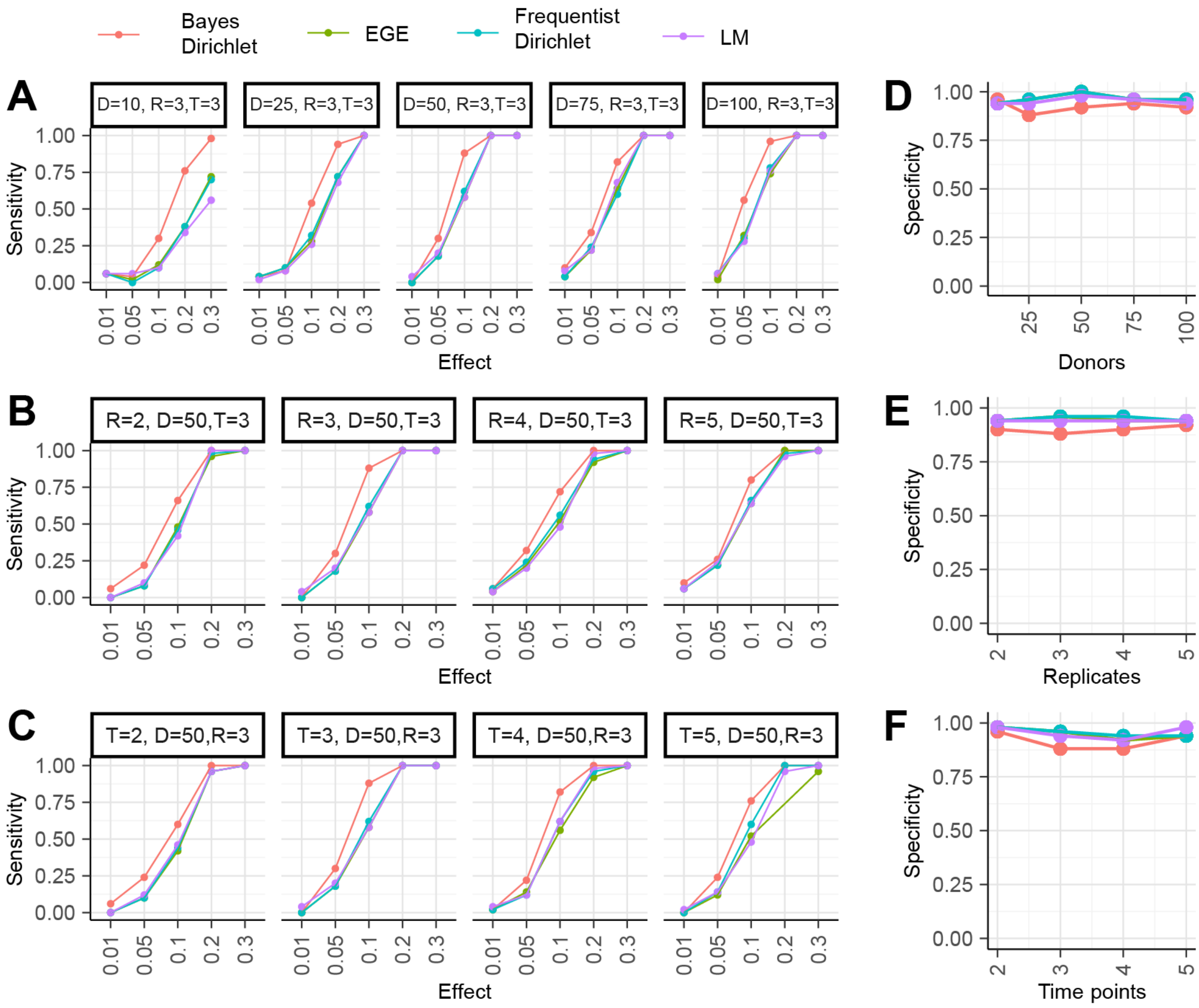


**Figure S2 Village design influences statistical power**
(A) Increasing the number of donors (D) has the greatest effect on sensitivity when both replicates (R) and sample time points (T) are held constant. (B) Townlet has increased sensitivity, especially when replication is low. (C) Increasing the number of sample time points does not necessarily increase power, likely due to donor takeover and underrepresentation towards the end of the experiment. (D-F) Specificity remains high when the (d) donor, (e) replicate, and (f) sample time point number are varied respectively.

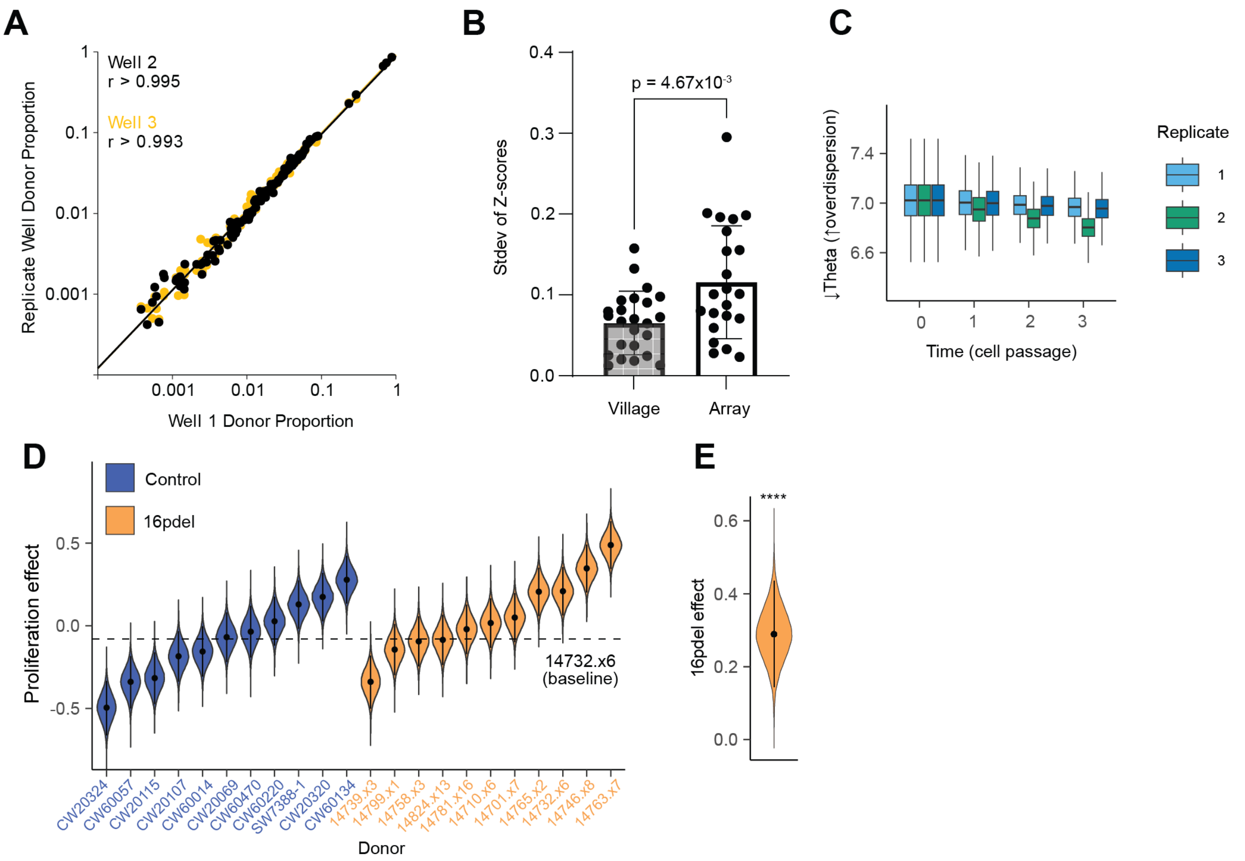


**Figure S3 Validation of 16pdel NPC hyperproliferation phenotype** (A) Donor proportions from 16p11.2 deletion village are highly correlated between replicates over all 6 time points (Well 2 vs Well 1, Pearson r = 0.995, Spearman r = 0.995; Well 3 vs Well 1, Pearson r = 0.993, Spearman r = 0.997) (B) Donor growth rates are normalized (for the village, within each well (n=3 wells). For the array, within each column (n=3 wells per donor)) to a mean of 0 and standard deviation of 1. Each data point represents the standard deviation across replicates of a donor's normalized growth rates (biological n = 23 donors; $p = 4.67 \cdot 10^{-3}$). (C) Technical variation is fairly consistent throughout the 16pdel experiment, but Replicate 2 has slightly more noise detected by Townlet. Decreasing values of theta equals more technical variation. (D) Townlet's individual total donor proliferation effect posteriors relative to the baseline donor (14732.x6). These were estimated from the 16pdel village while excluding the fastest deletion donor NFID_0176 (proportion data renormalized). Posterior means (points) and 95% credible intervals (errorbars) are shown. (E) Strong 16pdel effect on proliferation exists even when omitting the fastest deletion donor from the analysis ($lfsr = 2 \cdot 10^{-4}$).

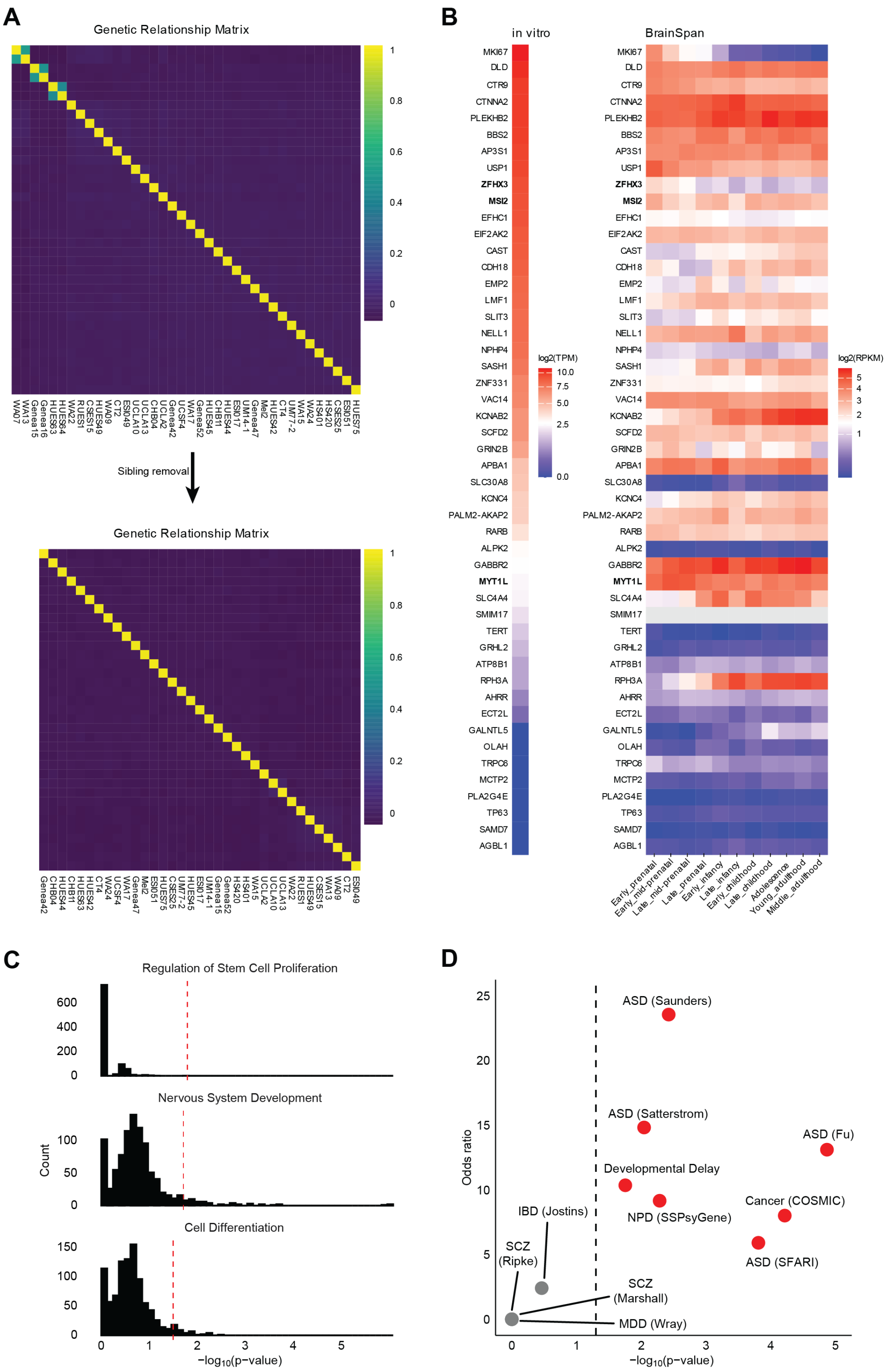


**Figure S4 Expression analysis of proliferation GWAS top loci** (A) Centered kinship coefficients (color viridis scale) estimated with GEMMA before and after removal of 3 siblings ($n = 38$ before removal, $n = 35$ after). (B) Left, expression data ($\log_2$ TPM, color heat scale) from bulk RNA-seq of *in vitro* SNaP model of human NPCs dataset[15]. Right, developmental expression ($\log_2$ RPKM, color heat scale) across 11 human brain stages ranging from 10 post-conception weeks to adulthood ($n = 49$ protein coding genes mapped from top 100 loci). Expression data obtained from BrainSpan[62]. (C) Distribution of GO term enrichment p-values from random shufflings of donor proliferation effects prior to GWAS ($n = 941$ permutations). Red dashed lines indicate observed significance in the unshuffled GWAS. (D) Summary of disease gene enrichment analysis. Red dots denote gene lists with significant overlap with mapped protein coding genes.

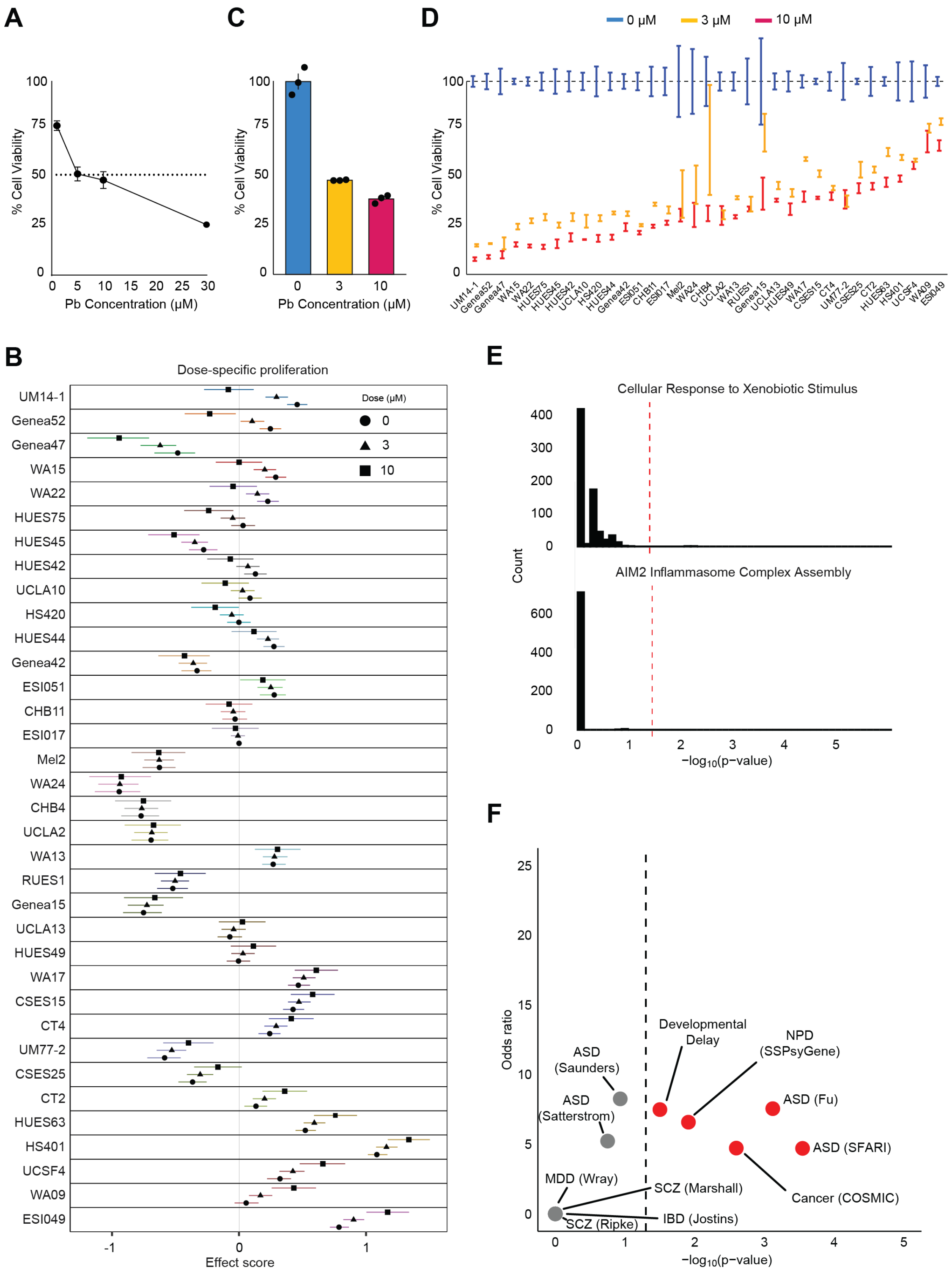


**Figure S5 Pb-induced reduction of NPC viability**

(A) Pb exposure reduces NPC viability in a dose-dependent manner. NPCs derived from the Genea52 ESC line were exposed daily to Pb (0, 1, 5, 10, 30 $\mu$M) for 7 days. Cell viability was quantified with CellTiter-Glo®2.0 and normalized to 0 $\mu$M (mean $\pm$ s.d., n = 3 wells per condition). (B) Townlet dose-specific total proliferation effect means with 95% credible interval error bars estimated per donor. (C) Cell viability of NPC village wells exposed daily to Pb (0, 1, 5, 10, 30 $\mu$M) for 7 days, estimated from brightfield-derived cell counts and normalized to the mean of 0 $\mu$M replicates (mean $\pm$ s.d., n = 3 wells per condition). (D) Donor-specific viabilities after Pb exposure. Donor viabilities were inferred by combining Census-seq donor proportions with replicate brightfield-derived cell counts. For each donor, inferred counts were normalized to the mean of 0 $\mu$M replicates (mean $\pm$ s.e.m., n = 3 wells per condition). (E) Distribution of GO term enrichment p-values from random shufflings of donor Pb effects prior to GWAS (n = 720 permutations). Red dashed lines indicate observed significance in the unshuffled GWAS. (F) Summary of disease gene enrichment analysis. Red dots denote gene lists with significant overlap with mapped protein coding genes.
